# *In Vitro* Proliferation and Long-Term Preservation of Functional Primary Rat Hepatocytes in Cell Fibers

**DOI:** 10.1101/2021.11.27.467621

**Authors:** Elsa Mazari-Arrighi, Teru Okitsu, Hiroki Teramae, Hoshimi Aoyagi, Mahiro Kiyosawa, Mariko Yano, Shoji Takeuchi

## Abstract

Primary hepatocytes are essential cellular resource for drug screening and medical transplantation. Since culture systems for them have already succeeded in reconstituting the biomimetic microenvironment, acquiring additional capabilities both to expand primary hepatocytes and to handle them easily would be expected as progress to the next stage. This paper describes a culture system for primary rat hepatocytes that is equipped with scalability and handleability relying on cell fiber technology. Cell fibers are cell-laden core-shell hydrogel microfibers; in the core regions, cells are embedded in extracellular matrix proteins, cultured three-dimensionally, and exposed to soluble growth factors in the culture medium through the hydrogel shells. By encapsulating primary rat hepatocytes within cell fibers, we first demonstrated they increase in number while keeping their viability and their hepatic specific functions for up to thirty days of subsequent culture. Then, we demonstrated the potency of the primary rat hepatocytes that proliferate in cell fibers not only as cell-based sensors to detect drugs that damage hepatic functions and hepatocellular processes but also as transplants to improve the plasma albumin concentrations of congenital analbuminemia. Therefore, our culture system could serve for innovating strategies and promising developments in applying primary hepatocytes to both pharmaceutical and medical fields.

## Introduction

Primary hepatocytes are currently used for fundamental studies as well as for pharmaceutical and medical applications including drug screening and hepatocyte transplantation^1–3^. Since the quality of hepatocytes is crucial for these applications, during the past years, culture systems for primary hepatocytes have made progress not only in long-term maintenance of their viability and their metabolic functions but also in allowing them to proliferate. In fact, these systems tend to enable primary hepatocytes *in vitro* to behave similarly to when they do *in vivo*; in the liver, functional hepatocytes retain their viability with a lifespan over three months^4^, while they proliferate by dividing until reaching the adequate cellular volume necessary for the liver to be restored when liver suffers from any change of losing its volume by injury, infection, or surgical procedure of partial hepatectomy^1^. Therefore, culture systems for primary hepatocytes have been developed to reproduce the microenvironment of hepatocytes in the liver, and three key factors are currently identified: i) three-dimensional culture^5,6^, ii) extracellular matrix (ECM) proteins^7,8^, and iii) soluble growth factors^9–11^. Recently, Rose et al. reported to succeed in allowing primary human hepatocytes to proliferate while keeping their metabolic functions for four weeks by embedding them in ECM of collagen where the hepatocytes formed spheroid-shaped clusters and are exposed to hepatocyte growth factor (HGF), epidermal growth factor (EGF), and insulin-transferrin selenium (ITS) within the medium^12^.

Once the culture system for primary hepatocytes has reconstituted this biomimetic microenvironment, the next key progress could be to acquire the capabilities of both scalability and handleability for the implementation into pharmaceutical and medical fields, because drug screening and hepatocyte transplantation require not only large numbers of hepatocytes^13–17^ but also easy-to-handle and transferable constructs in which hepatocytes are cultured^17,18^. However, few investigations have been performed to yield such a scalable and handleable culture system for primary hepatocytes; although core-shell hydrogel microcapsules might be good strategy^19–21^, they still lack the evidence to show the potential of their practical use through providing primary hepatocytes with biomimetic culture environment accompanied with both scalability and handleability.

Cell fiber is a useful system to culture cells three-dimensionally for a long term; in cell fibers, the cells can proliferate, migrate, and connect with each other to form a functional cellular tissue^22^. Cell fibers are cell-laden core-shell hydrogel microfibers and they are generated by using a double-coaxial laminar-flow microfluidic device; the core contains both cells and ECM proteins while the shell consists of alginate hydrogel. All the cells in the core can easily access oxygen and nutrients in the culture medium since the thickness of the cell fiber is kept a few hundreds of micrometres over its entire length. Cell fibers have been shown to serve for three-dimensional tissue formation of various types of cells including cardiomyocytes, vascular endothelial cells, nerve cells, smooth muscle cells and adipocytes^22–25^. The advantage of cell fiber is to be mass-producible and scalable; a cell fiber with long (meter-order) length can be fabricated in a short time (a few minutes) and cell fibers have allowed pluripotent stem cells to increase in number while keeping their original characters^24^. The other advantage of cell fiber is to be handleable; cell fibers encapsulating pancreatic islet cells have been stably and reproducibly transplanted to diabetic mice and even retrieved from these animals^22^.

In this study, we hypothesize that cell fibers can be used to develop a culture system for primary hepatocytes that is equipped with scalability and handleability, in addition to the already available culture biomimetic environment that allows primary hepatocytes both to proliferate and to maintain their viability as well as metabolic functions. Here, we first describe our encapsulation technique of primary rat hepatocytes into core-shell hydrogel microfibers to form cell fibers. Secondly, we evaluate whether primary rat hepatocytes cultured in cell fibers can proliferate and can subsequently preserve their viability and their functionality for up to 30 days. We then demonstrate pharmaceutical relevance of the primary rat hepatocytes cultured in cell fibers for screening drugs that might damage hepatic functions and hepatocellular processes such as proliferation. Finally, to show scalability and handleability of our culture system for the treatment of the diseases due to hepatocellular malfunction, we assess the potency of the primary rat hepatocytes cultured in cell fibers as transplants to improve the plasma albumin concentrations in analbumenic rats.

## Results

### Applying cell fiber technology to encapsulate primary rat hepatocytes within cell fibers

To embed primary rat hepatocytes in ECM proteins for providing them with three-dimensional culture environment, we used cell fiber technology that we have developed^22^. First, we evaluated the impact of the initial cell seeding density of primary rat hepatocytes both on how they would occupy the space of the core region of cell fibers and on how they would behave there afterwards. For this evaluation, core-shell hydrogel microfibers were fabricated to encapsulate primary rat hepatocytes through a double-coaxial laminar-flow microfluidic device (**Figure 1a**) by applying the ECM conditions that we optimized beforehand [a mixture of type I collagen (Native collagen) and Matrigel (**Figure S1**)] and by using the following three different initial cell seeding densities of primary rat hepatocytes: 2.5×10^7^, 5×10^7^, and 9×10^7^ cells mL^-1^. Through microscopic observations just after cell fiber fabrication, we found that, when the initial cell seeding density was 9×10^7^ cells mL^-1^, the encapsulated cells nearly fully occupied the core regions of the cell fibers, and the space of the core regions occupied by the cells decreased normally as the initial cell seeding density goes down (**Figure 1b, d, f**). After 48 hours of culture without any stimulation of proliferation, we found in their core regions that the encapsulated cells self-assembled into different cellular clusters in terms of shape and size, depending on each initial cell seeding density; at 9×10^7^ cells mL^-1^, primary rat hepatocytes formed fiber-shaped clusters; at 5×10^7^ cells mL^-1^, they formed rod-shaped clusters; at 2.5×10^7^ cells mL^-1^, they formed mostly spheroid-shaped clusters (**Figure 1c, e, g**). These results indicate that primary rat hepatocytes cultured in cell fibers gather spontaneously to form cellular clusters during 48 hours after being encapsulated, and that the shape as well as the size of resulting cellular clusters would vary based on the initial cell density embedded in the core region; within a certain window of the initial cell density, the higher it is, the more likely to be fiber-shaped the cellular clusters would be; in contrast, the lower it is, the more likely to be spheroid-shaped.

**Figure 1.**
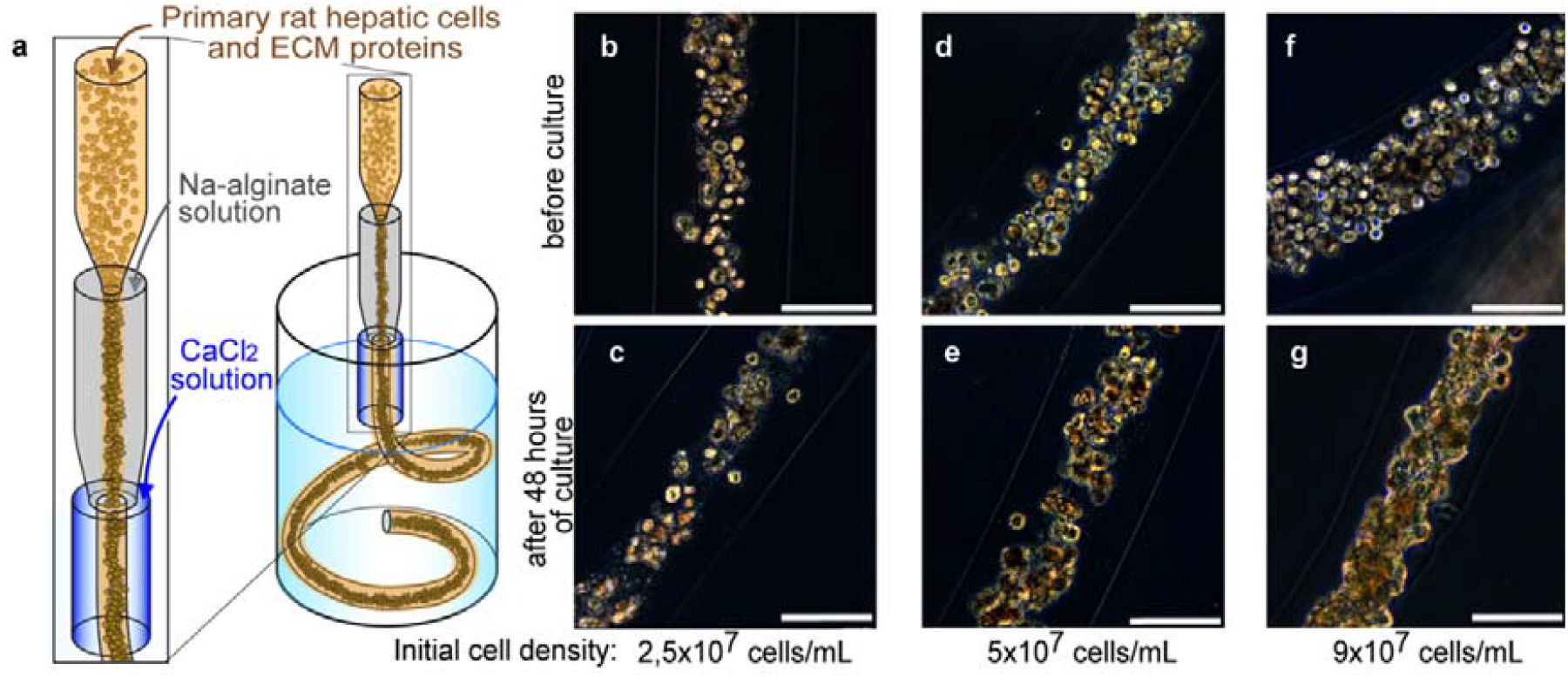
Encapsulating primary rat hepatocytes within core-shell hydrogel microfibers by applying cell fiber technology. (**a**) Schematic drawing of the fabrication of core-shell hydrogel microfibers encapsulating freshly isolated rat hepatocytes through a double co-axial microfluidic device. (**b-g**) Representative dark-field images (n=12 cell fibers for each group) of primary rat hepatocytes encapsulated in cell fibers before culture and after 48 hours of culture in three experimental groups possessing different initial cell seeding densities: 2,5×10^7^ cells mL^-1^, 5×10^7^ cells mL^-1^, and 9×10^7^ cells mL^-1^. Scale bars; 100 µm.

### *In vitro* proliferation and long-term maintenance of functional primary rat hepatocytes in cell fibers

We hypothesized that cell fiber could create an appropriate microenvironment for soluble factor stimulation to trigger the proliferation of encapsulated primary rat hepatocytes. Before testing this hypothesis, as described in literatures^9–12^, we prepared the already identified soluble factors including HGF and EGF to stimulate the proliferation. Eventually, we decided to use the conditioned medium collected from 3T3 cells (3T3CM) that had been cultured in cell fibers for 5 days, because we confirmed that the amount of HGF measured in the 3T3CM is the highest after 5 days of culture (**Figure S2g**), and also that this 3T3CM is comparable with the medium supplemented with not only recombinant mouse HGF but also recombinant mouse EGF (**Figure S3m-n**).

Then, to test our hypothesis, primary rat hepatocytes were encapsulated into core-shell hydrogel microfibers at the initial cell seeding density of 2.5×10^7^ cells mL^-1^; this value was determined based on the findings of our previous experiments showing that such initial cell seeding density secures space for the cultured primary rat hepatocytes to increase in number within cell fibers (**Figure 1b, d**). After 2 days of culture, cell fibers that encapsulate primary rat hepatocytes at 2.5×10^7^ cells mL^-1^ were randomly divided into two groups: 1) in the experimental group, cell fibers started to be cultured in media containing 50% of 3T3CM, 2) in the control group, cell fibers were cultured in the same medium as the one used for the first 2 days of culture. Subsequently, we assessed the morphologies of primary rat hepatocytes cultured in those cell fibers, we evaluated various characteristics including hepatocyte-specific functions: (i) cell number, (ii) viability, (iii) albumin secretion, (iv) urea synthesis, (v) CYP1A1 enzyme activity, and we compared these characteristics between the experimental and the control groups for up to 30 days of culture. Regarding microscopic morphologies, as we expected, primary rat hepatocytes cultured in cell fibers for up to 2 days gather spontaneously into spheroid-shaped clusters that were scattered in the core regions of cell fibers (**Figure 2b, f**). After 4 days of culture *via* the random division, we observed that primary rat hepatocytes occupied almost the space of the core regions and formed fiber-shaped aggregates in the experimental group, while they kept spheroid-shaped clusters being sparsely scattered in the control group (**Figure 2c, g**). Furthermore, we found in the experimental group that the cell number increased to be 2.4±0.7 times higher after 4 days of culture than after 2 days of culture and that the viability of 45.8±2.0% was maintained up to 30 days of culture. In contrast, in the control group that neither the cell number increased, nor the viability maintained (**Figure 2i-j**). We also found that primary rat hepatocytes maintained their hepatocyte-specific functions (albumin secretion, urea synthesis, CYP1A1 enzyme activity) up to 30 days of culture in the experimental group, but not in the control group (**Figure 2k-m**).

**Figure 2.**
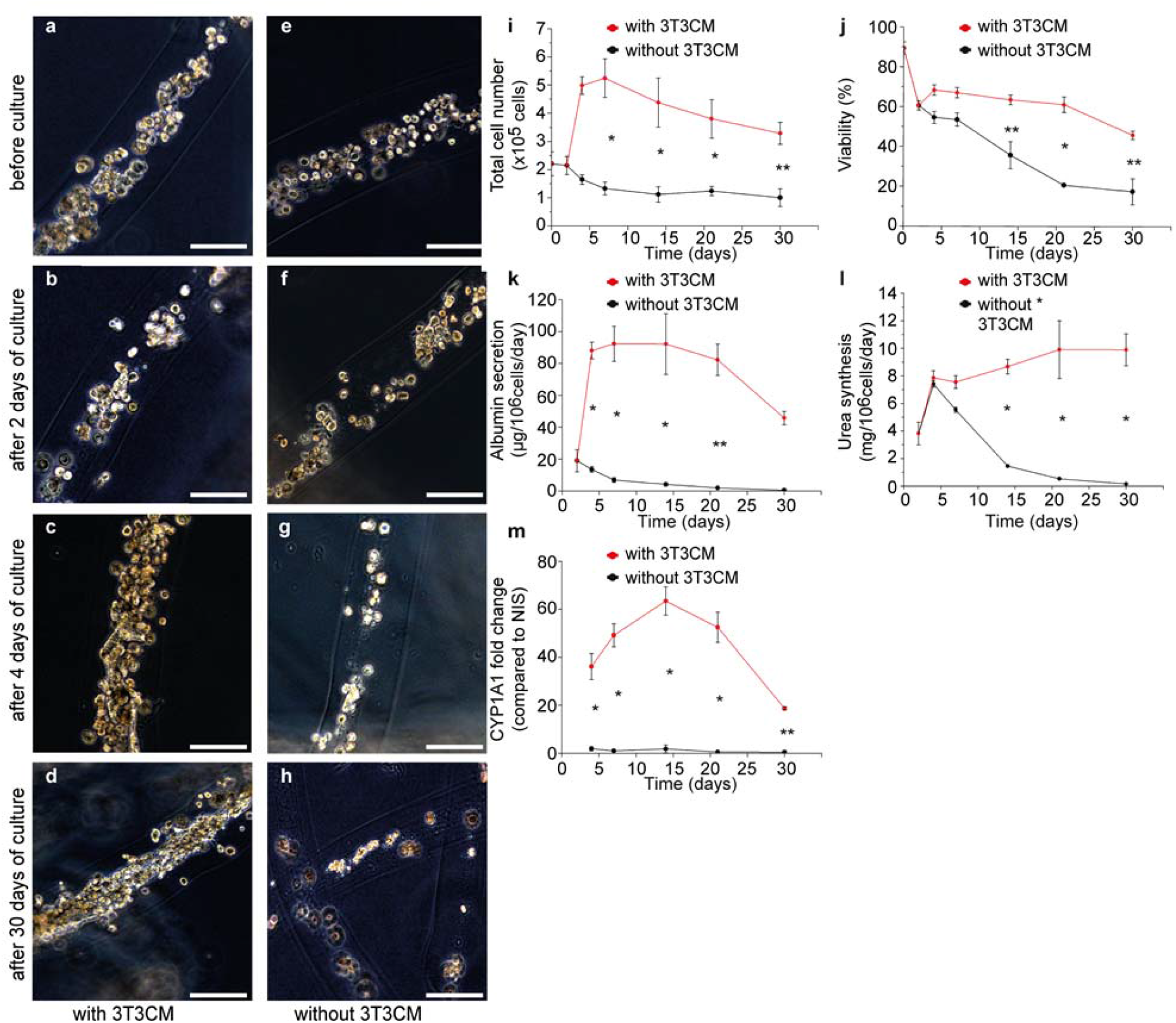
Culture of primary rat hepatocytes in cell fibers for up to 30 days either with or without using 3T3CM. (**a-h**) Representative dark-field images of primary rat hepatocytes encapsulated in cell fibers (n=12 for each group) at the initial cell seeding density of 2.5×10^7^ cells mL^-1^ before culture, after 2 days, after 4 days, and 30 days of culture in two experimental groups: (**a-d**) in one group 3T3CM was used after 2 days of culture, (**e-h**) while in the other group 3T3CM was not used. Scale bars; 100 µm. (**i-m**) Time course of the characteristics including hepatocyte-specific functions of primary rat hepatocytes cultured in cell fibers (n=4 per data point) for over 30 days of culture either with or without using 3T3 CM: (**i**) total cell number, (**j**) viability, (**k**) albumin secretion, (**l**) urea synthesis, (**m**) CYP1A1 enzyme activity. Error bars denote mean ± SD. *P < 0.01 and **P < 0.05; Student’s t-test at each time point.

Thereafter, we confirmed whether the cells in the experimental group, which have proliferated under proliferation stimulation in cell fibers, were hepatocytes or not. We then analysed the expression of a specific marker of hepatocyte, ASGPR-1, on the encapsulated cells in cell fibers using flow cytometry^26^ and we also attempted to detect albumin in those cells by using immunohistochemistry. We measured that the number of ASGPR-1 expressing cells were 219% higher after 7 days of culture than before stimulation, and that the number of ASGPR-1 expressing cells were still 104% higher after 30 days of culture than before stimulation (**Figure 3a-d, Table S1**). We also found that almost all encapsulated cells that have been cultured in the cell fibers for 7 days were albumin positive and they seemed to connect with each other (**Figure 3e-f**).

**Figure 3.**
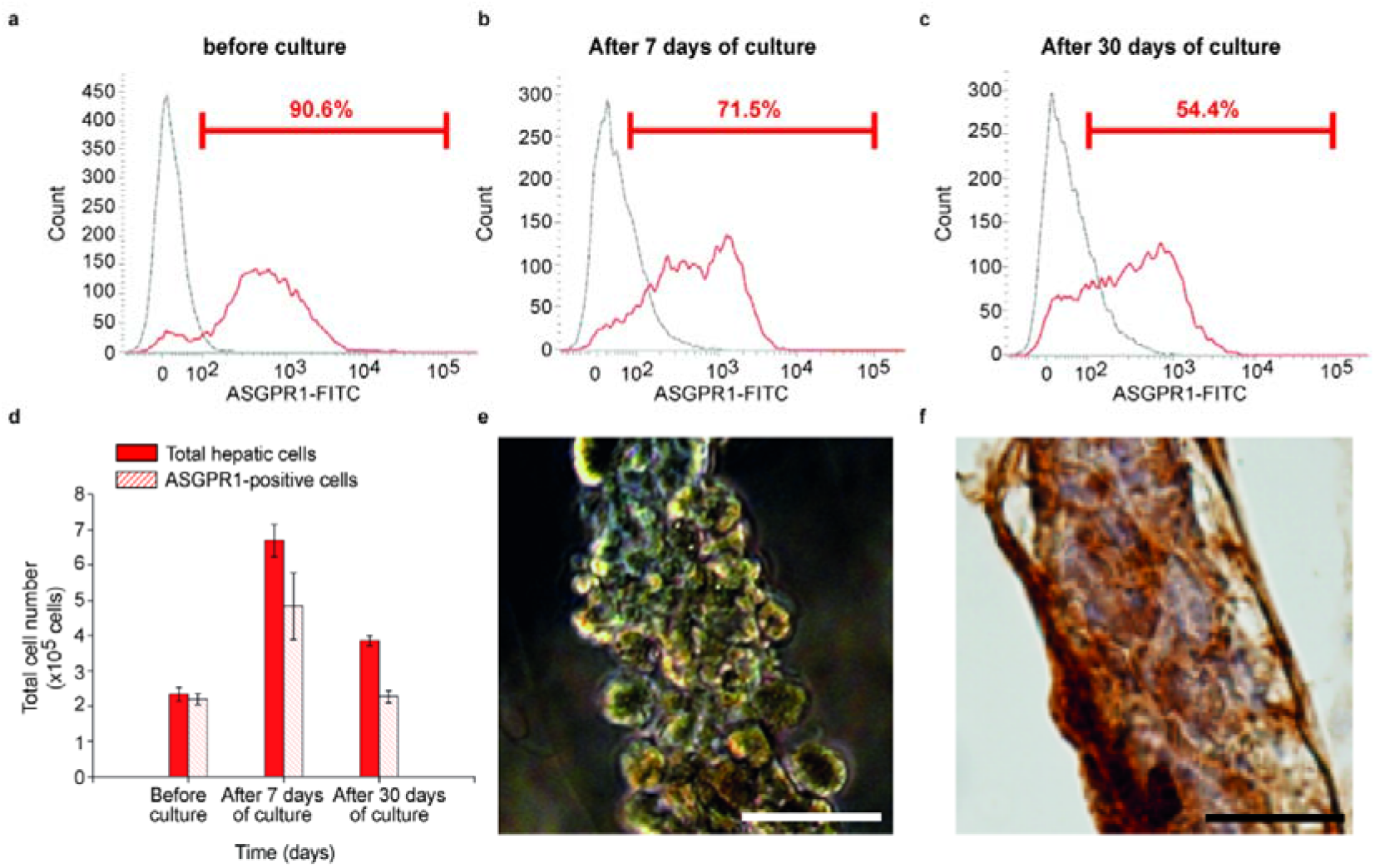
Hepatocyte detection in primary rat hepatic cells cultured in cell fibers using 3T3CM. (**a**-**c**) Representative FACS histograms (of 3 samples at least for each culture duration) showing the surface marker of asialoglycoprotein receptor-1 (ASGPR-1) on the primary rat hepatocytes (**a**) before culture, (**b**) after 7 days, (**c**) and 30 days of culture in cell fibers using 3T3CM. Red lines represent samples that were treated using both of a mouse-anti-human ASGPR1 antibody and a FITC-labelled goat anti-mouse antibody, while grey lines represent samples that were treated only with a FITC-labelled goat anti-mouse antibody. Red numbers in FACS histograms represent the percentage of cells that are positive for ASGPR-1. (**d**) Total cell number and ASGPR-1 positive cell number in primary rat hepatocytes before culture, after 7 days, and 30 days of culture (n=3 at least per each time point). (**e**) Representative dark-field image (n=3) and (**f**) representative image of immunohistochemical analysis for albumin localization (n=3) of primary rat hepatocytes that were cultured for 7 days in cell fibers using 3T3CM. Scale bars; 50 µm.

Taken together, these results clearly show that primary rat hepatocytes can survive for a long term while keeping their functions in cell fibers only when they fill the core space by increasing in number within 4 days of culture. These results also indicate that primary rat hepatocytes should connect with each other and form cellular aggregates in order to acquire three-dimensional culture environment; for primary rat hepatocytes to connect with ECM proteins is not enough to survive in cell fibers for a long term.

### *In vitro* detection of drug hepatotoxicity using primary rat hepatocytes in cell fibers

To explore the possible application of cell fibers encapsulating primary hepatocytes in the field of drug screening, we evaluated potential of cell fibers for *in vitro* detection of hepatotoxicity of drugs. For this evaluation, we attempted to obtain the respective concentrations of a 50% inhibitory effect (IC50 values) for two kinds of well-known hepatotoxic compounds including acetaminophen and diclofenac. First, primary rat hepatocytes were encapsulated into core-shell hydrogel microfibers at the initial cell seeding density of 2.5×10^7^ cells mL^-1^ and were cultured in the media supplemented with 3T3CM. Secondly, the concentration-reaction curves for these compounds were obtained through the assays of cellular viability, albumin secretion, and urea synthesis performed at different time points: 4 days, 7 days, 14 days, and 30 days after the start of culture. Thirdly, based upon these concentration-reaction curves (**Figures S4-S7**), the corresponding IC50 values were estimated. Moreover, we attempted to compare these IC50 values with those obtained using the other culture system where primary rat hepatocytes were cultured on collagen-coated 24-well plates. We found that the IC50 values for both acetaminophen and diclofenac can be estimated up to 30 days of culture when using cell fibers, and that these estimated IC50 values for each compound are both reproducible in individual cell fibers as well as stable over time regardless of these three assay methods, while the IC50 values for either of the two compounds become unable to be estimated after more than 7 days of culture when using collagen-coated 24-well plates (**Table 1**). We also found that the IC50 values for both the compounds obtained using cell fibers are in good accordance with the IC50 values reported previously in the literature (**Tables S2**-**S3**)^27–30^. These findings indicate that the culture system using cell fibers encapsulating primary rat hepatocytes could be useful to estimate drug hepatotoxicity in the field of drug screening and might even improve the reliability of the estimation because of the reproducibility and the chronological stability of the primary rat hepatocytes cultured in the cell fibers (**Table S4**).

**Table 1.**
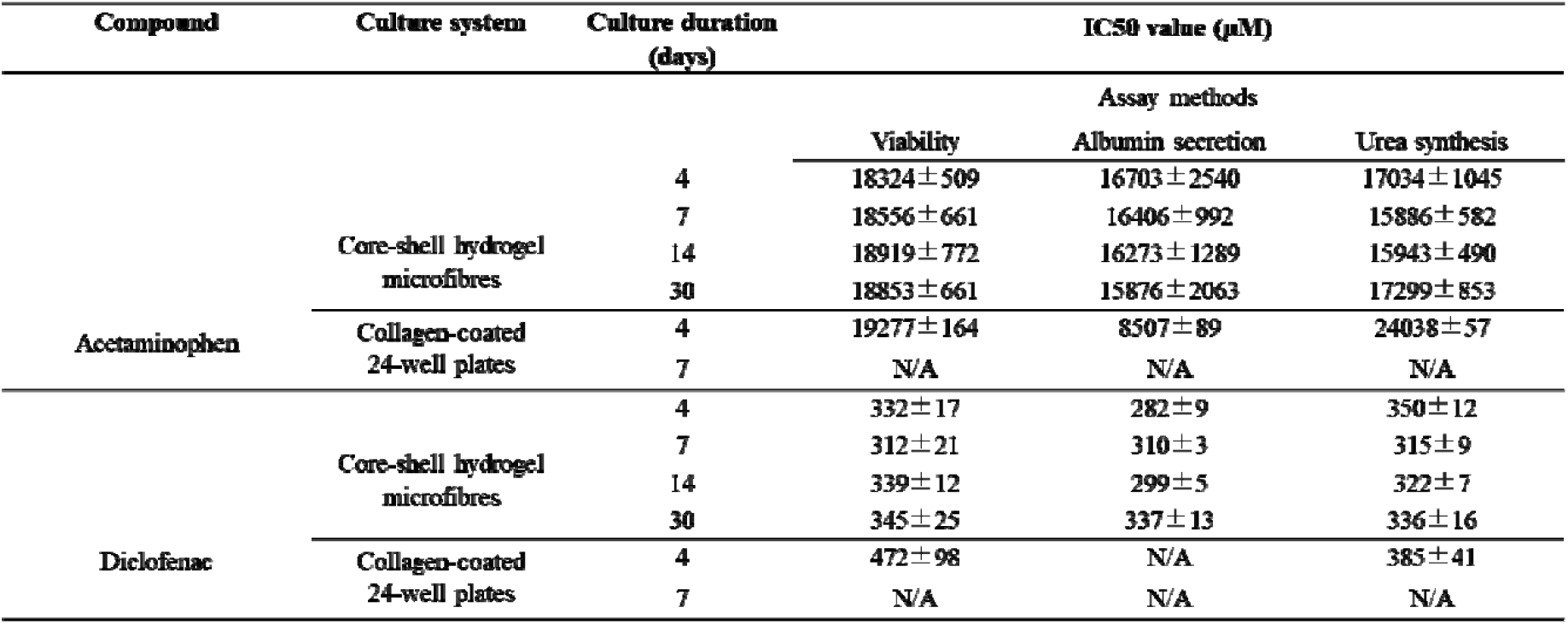
50% inhibitory effect (IC50 values) for acetaminophen and diclofenac relating to viability, albumin secretion, and urea synthesis of primary rat hepatocytes cultured either in cell fibers for 4 days, 7 days, 14 days, and 30 days, or in collagen-coated 24-well plates for 4 days and 7 days. Data are presented as the mean ± standard deviation of at least four independent cell fibers and four collagen-coated wells from collagen-coated 24-well plates.

| Compound | Culture system | Culture duration (days) | IC50 value (μM) | | |
| --- | --- | --- | --- | --- | --- |
|  |  |  | Assay methods |  |  |
|  |  |  | Viability | Albumin secretion | Urea synthesis |
| Acetaminophen | Core-shell hydrogel microfibres | 4 | 18324 ± 509 | 16703 ± 2540 | 17034 ± 1045 |
| | | 7 | 18556 ± 661 | 16406 ± 992 | 15886 ± 582 |
| | | 14 | 18919 ± 772 | 16273 ± 1289 | 15943 ± 490 |
| | | 30 | 18853 ± 661 | 15876 ± 2063 | 17299 ± 853 |
| | Collagen-coated 24-well plates | 4 | 19277 ± 164 | 8507 ± 89 | 24038 ± 57 |
|  |  | 7 | N/A | N/A | N/A |
| Diclofenac | Core-shell hydrogel microfibres | 4 | 332 ± 17 | 282 ± 9 | 350 ± 12 |
| | | 7 | 312 ± 21 | 310 ± 3 | 315 ± 9 |
| | | 14 | 339 ± 12 | 299 ± 5 | 322 ± 7 |
| | | 30 | 345 ± 25 | 337 ± 13 | 336 ± 16 |
| | Collagen-coated 24-well plates | 4 | 472 ± 98 | N/A | 385 ± 41 |
|  |  | 7 | N/A | N/A | N/A |

### *In vitro* detection of drug inhibition of hepatic regeneration using primary rat hepatocytes in cell fibers

Since we have revealed that primary rat hepatocytes can proliferate in cell fibers while keeping their hepatocyte-specific functions, we hypothesized that these cell fibers could serve for screening drugs that inhibit hepatic regeneration. To test this hypothesis, we adopted retrorsine, a compound that is standardly used to inhibit rat hepatocyte regeneration *in vivo*^31,32^ and we attempted to measure its IC50 value by obtaining its concentration-reaction curve. Practically, core-shell hydrogel microfibers were fabricated to encapsulate primary rat hepatocytes at the initial cell seeding density of 2.5×10^7^ cells mL^-1^. Then, the fabricated cell fibers were randomly divided into four groups; 1) in the control group, the cell fibers were cultured in media containing 3T3CM after 2 days of culture, 2) in the three experimental groups, the cell fibers were cultured under the same condition as the control group except being exposed to three different concentrations of retrorsine (2.5 mg mL^-1^, 5 mg mL^-1^, 10 mg mL^-1^) for 24 hours after 2 days of culture. The proliferation ratio of primary hepatocytes in the experimental groups were calculated, and the concentration-reaction curve for retrorsine was obtained (**Figure 4**). Using the curve, we found that the IC50 value for retrorsine is estimated to be 3.1 ± 0.1 mg mL^-1^ (∼8.82 ± 0.3 mM).

**Figure 4.**
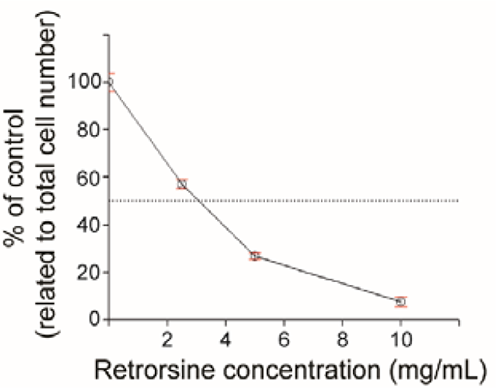
Retrorsine concentration-reaction curve relating to inhibition of primary rat hepatocyte proliferation in cell fibers *in vitro*. The effect of 24-hour retrorsine treatment on inhibition of primary rat hepatocyte proliferation in cell fibers was evaluated using three different concentrations (2.5 mg mL^-1^, 5 mg mL^-1^, 10 mg mL^-1^) of retrorsine (n=5 per data point). Error bars denote mean ± SD.

Subsequently, we evaluated whether this inhibitory effect of retrorsine on hepatocyte proliferation is reversible or irreversible. For that, we first fabricated cell fibers encapsulating primary rat hepatocytes at the initial cell seeding density of 2.5×10^7^ cells mL^-1^. Then, after 2 days of culture, we randomly divided them into two groups; in the control group, the cell fibers were cultured in media containing 3T3CM for up to subsequent 5 days of additional culture; in the experimental group, the cell fibers were cultured under the same condition as the control group except being exposed to retrorsine at a concentration of 10 mg mL^-1^ during 24 hours from day 2 of culture to day 3 of culture. Subsequently, we compared these two groups over time from viewpoints both of microscopical morphology and cell number. We found microscopically that primary rat hepatocytes in the experimental group proliferated after the removal of retrorsine and became fiber-shaped aggregates similar to those in the control group after 7 days of culture (**Figure 5a-b)**. We also found that the number of the cells in cell fibers of the experimental group did not change during the exposure to retrorsine, started to increase just after the removal of retrorsine in the culture medium, and became comparable to those in the control group after 7 days of culture (**Figure 5c)**. These results indicate that cell fibers where primary rat hepatocytes can proliferate could be useful for *in vitro* detection of drugs that inhibit hepatic regeneration and could be efficient to evaluate whether this inhibitory effect on hepatic regeneration is temporary or permanent.

**Figure 5.**
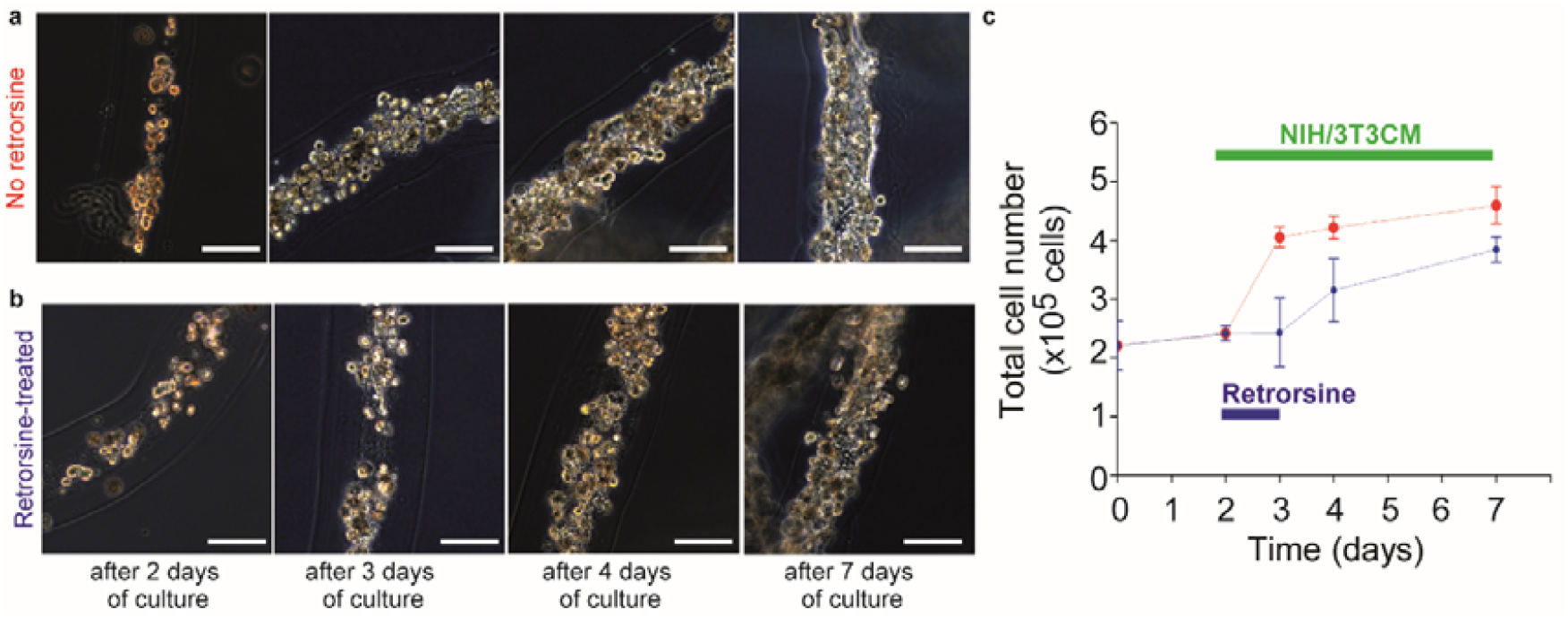
Proliferation of primary rat hepatocytes after 24-hour treatment of retrorsine in cell fibers. (**a**-**b**) Representative dark-field images of primary rat hepatocytes cultured in cell fibers (n= 5 for each group) for 2 days, 3 days, 4 days, and 7 days using 3T3CM. Cell fibers encapsulating primary hepatocytes were divided into two groups: (**a**) in one group the hepatocytes were not treated with retrorsine, (**b**) while in the other group they were treated with retrorsine for 24-hours after 2 days of culture. Scale bars; 100 µm. (**c**) Time course of total cell number of primary rat hepatocytes cultured in cell fibers using 3T3CM either (**c**) with or (**d**) without 24-hour retrorsine treatment after 2 days of culture (n=5 per data point). Error bars denote mean ± SD.

### *In vivo* albumin secretion function of primary rat hepatocytes that have proliferated and cultured in cell fibers

We investigated further applicability of cell fibers encapsulating primary hepatocytes to the field of medical transplantation for the treatment of a metabolic disease, especially analbuminemia^33^. For this investigation, we prepared two types of grafts and four groups of at least four Nagase analbuminemia rats (NARs). To prepare two types of grafts, core-shell hydrogel microfibers were fabricated to encapsulate primary rat hepatocytes at the initial cell seeding density of 2.5×10^7^ cells mL^-1^ and, 2 days after fabrication, the cell fibers were randomly divided into two groups. In one group, the cell fibers were cultured for 5 more days in media supplemented with 3T3CM resulting in proliferation of the encapsulated primary rat hepatocytes, this graft type is called “cell fiber grafts with 3T3CM”. For the other graft type, cell fibers were cultured for 5 more days without being exposed to 3T3CM resulting in no proliferation of the encapsulated primary rat hepatocytes, which is called: “cell fiber grafts without 3T3CM”. Before transplantation experiments, NARs were randomly assigned into four groups: one experimental group and three control groups. In the experimental group, the NARs received the cell fiber grafts with 3T3CM. In the first control group, the NARs received the cell fiber grafts without 3T3CM. For experimental group, prior to transplantation, we evaluated the number of cells within the cell fiber grafts with 3T3CM and we transplanted the cell fibers containing approximately 2.0×10^7^ cells in total by putting them into each intra-mesenteric space of at least four NARs (**Figure 6a-c, Figure S8**)^34^. In parallel, for the first control group, we transplanted cell fiber grafts with the same total length as those transplanted in the experimental group, also by putting them into the same site of four NARs. All the NARs in both these groups received daily injections of tacrolimus, an immunosuppressive drug also named FK-506, after transplantation. In the second and the third control groups, the NARs received no transplant, but the former received daily injections of FK-506, and the latter received no treatment.

**Figure 6.**
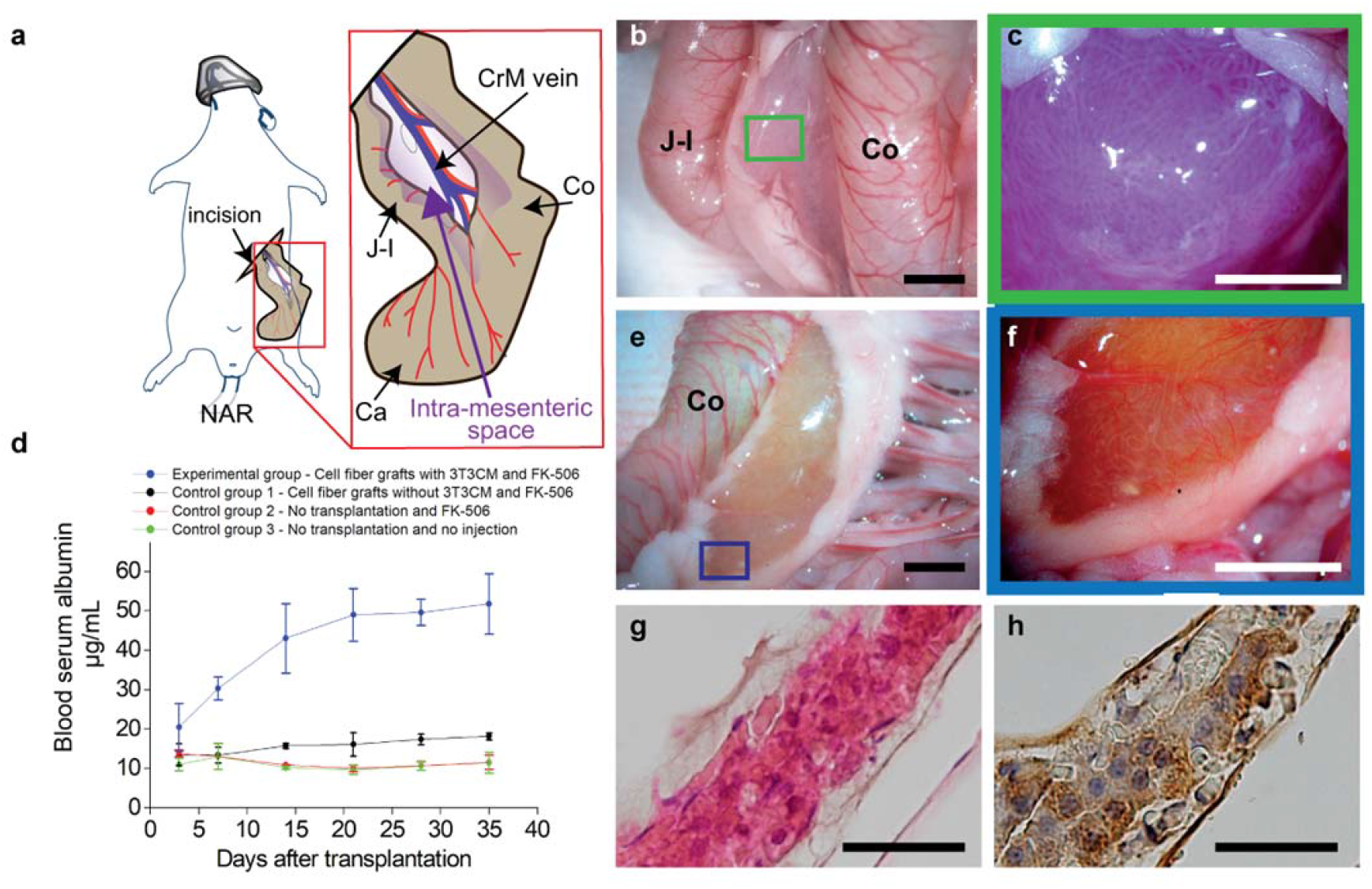
Transplantation of cell fibers encapsulating primary rat hepatocytes to analbumenic rats. (a) Schematic drawing of intra-mesenteric space as the transplantation site for cell fibers encapsulating primary rat hepatocytes. (**b-c**) Representative photo images (n=5) with (**b**) low and (**c**) high magnifications of cell fibers in intra-mesenteric space immediately after transplantation. Scale bars; 5 mm in **(b)**; 2 mm in **(c)**. (**d**) Time course of plasma albumin concentrations of the analbumenic rats (NARs) in 4 groups; one group received transplants of the cell fibers cultured with 3T3 CM and daily injections of immunosuppressive drug FK-506, one group received transplants of the cell fibers cultured without 3T3 CM and the daily injections, one group received no transplants but only daily injections, one group receive neither transplants nor daily injections (n=4 at least for each group). Error bars denote mean ± SD. *P < 0.01 and **P < 0.05; Student’s t-test at each time point. (**e-f**) Representative photo images (n=4) with (**e**) low and (**f**) high magnifications of cell fibers in intra-mesenteric space 35 days after transplantation. Scale bars; 5 mm in (**e**); 2 mm in (**f**). (**g-h**) Representative images of histological analysis (**g**) using haematoxylin and eosin staining, and (**h**) of immunohistochemical analysis for albumin localization of primary rat hepatocytes in cell fibers retrieved 35 days after transplantation (n=4 rats for each analysis). Scale bars; 200 µm. CrM vein, Crania Mesenteric vein; J-I, Jejuno-Ileum; Ca, Caecum; Co, Colon.

We then found that plasma albumin concentrations of NARs increased over time up to 35 days after transplantation in the experimental group and that the amount of plasma albumin concentration was approximately three to five times higher compared with all the control groups (**Figure 6d**). Furthermore, thirty-five days after transplantation, we performed second-look analysis for NARs of the experimental group, and we observed that the transplanted cell fibers containing primary rat hepatocytes were microscopically recognizable in the intra-mesenteric spaces of all NARs (**Figure 6f, Figure S9)**. In addition, histological analysis revealed that the cells in the cell fibers are morphologically intact (**Figure 6g**) and that almost all these cells were stained with albumin (**Figure 6h**). These results clearly show that primary rat hepatocytes that had *in vitro* proliferated and were cultured in cell fibers could fulfil their ability to secrete significant level of albumin *in vivo*, and that these cell fibers could be easily handled as transplants to treat analbuminemia. Taken together, these results confirm that the cell fibers encapsulating primary rat hepatocytes are equipped with handleable and scalable characteristics that would be essential in the context of transplantation of primary hepatocytes.

## Discussion

In the present study, we have developed, using cell fiber technology, a culture system equipped with scalability and handleability for primary rat hepatocytes that allows them both to proliferate and to retain their viability as well as metabolic functions for a long term within cell fibers. The evidence showing that our culture system allows primary rat hepatocytes both to proliferate and to maintain their viability as well as metabolic functions for a long term is that primary rat hepatocytes encapsulated in cell fibers increased in number under stimulation of specific environmental factors (**Figure 3 and Figure S3**); subsequently over culture time, these primary hepatocytes that have proliferated maintained their number and viability for at least 30 days while keeping their metabolic functions for the same period of time (**Figure 2**). Furthermore, the evidence showing that our culture system acquired capabilities of both scalability and handleability is that the primary rat hepatocytes encapsulated within cell fibers increased in number while keeping not only their sensitivity sufficient to detect the toxic effect of well-known hepatotoxic compounds such as acetaminophen and diclofenac (**Table 1 and Figures S4-S7**), but also their ability to secrete albumin in adequate amount to be observed in the blood of recipient rats with analbuminemia after being transplanted into NARs (**Figure 6**). As a matter of fact, all these experimental operations were performed with stability and reproducibility certainly thanks to the easiness with which the cell fibers are able to be fabricated, handled, and transferred.

In addition, we have shown a potency of our culture system to screen candidate compounds with either promoting or inhibitory effect on hepatic regeneration. This potency is based on the evidence that primary rat hepatocytes in cell fibers under proliferation stimulation lacked increasing in number while they are exposed to retrorsine, a well-known hepatotoxic compound to disrupt the division of rat hepatocytes; however, after ceasing exposure to retrorsine, the primary rat hepatocytes started proliferating and eventually formed fiber-shaped cellular aggregates (**Figure 5**). We also believe that this potency of our culture system could expand on reproducing a phenomenon seen *in vivo*: in the retrorsine-treated rats, the regeneration of their livers delayed comparing with the non-treated control rats after they underwent partial hepatectomy that exerts strong proliferative stimulus on the liver through activating several pathways including raising growth factors like HGF and EGF^35–37^ but the retrorsine-treated rats eventually regenerated their liver mass completely^32^. Additionally, as a noteworthy fact showing that our culture system provides handleable cell fibers, stopping exposure to retrorsine was experimentally done by transferring the cell fibers from a medium supplemented with retrorsine to another medium without retrorsine; these procedures could probably allow the on-off switch of the effect of retrorsine to get sharpened while minimizing the damage to the primary rat hepatocytes encapsulated within cell fibers.

Moreover, in this study, we demonstrated that our culture system within which primary hepatocytes have proliferated are potent to be applied for the field of medical transplantation by confirming that, after increasing in number within cell fibers, the primary rat hepatocytes improved plasma albumin concentrations in analbuminemia rats (**Figure 6**). We believe that cell fibers encapsulating primary hepatocytes possess two advantageous key aspects in this field. As qualitative aspect, the primary hepatocytes encapsulated in cell fibers connect with each other through ECM proteins, and are expected to fulfil better hepatocyte-specific functions than the one of the same number of dispersed primary hepatocytes^31,33,38–40^. Furthermore, as a quantitative aspect, our cell fibers allow primary hepatocytes to proliferate and can expand the supply of primary hepatocytes, potentially resulting in improvement in transplant preparation especially when isolated primary hepatocytes are short in amount.

Taken together, here we have shown that cell fibers could serve to establish a culture system equipped with scalability and handleability for primary rat hepatocytes that are cultured under biomimetic microenvironment for a long-term period; within the cell fibers, primary rat hepatocytes have proliferated and subsequently maintained their viability as well as their specific hepatic functions for up to 30 days. Adding on to these features, scalability and handleability of cell fibers have served for realizing operation procedures necessary for stability and reproducibility required in drug screening and medical transplantation. We believe that these features of cell fibers should lead to innovating strategies and promising developments in applying primary hepatocytes to both pharmaceutical and medical fields.

Finally, regarding the species origin of the primary hepatocytes, in the present study, we used laboratory rats under the consideration that they would be preferred for the initial development stage of our culture system for primary hepatocytes, because laboratory small animals could supply primary hepatocytes possessing stable quality through a whole series of experiments, and rats can provide a sufficient quantity of primary hepatocytes for each experiment of this stage. As the next step toward establishment of our culture system for primary hepatocytes, we are planning to use primary human hepatocytes. Although there is interspecies difference in the function of primary hepatocytes between rats and human^41^, we are quite optimistic about the development of such culture system using human primary hepatocytes, since Rose et. al. has paved the way for this kind of development using spheroids^12^, and we also have already succeeded in developing cell fibers for different types of human-derived cells23–25,42

## Material and methods

### Hepatocyte isolation

All animal experiments in this study were approved in advance by the University of Tokyo (approved protocol ID: 26–12) and were conducted in accordance with the rules of its Institutional Animal Care and Use Committee. Primary hepatocytes were isolated from six- to nine-week-old male Sprague-Dawley rats (Japan SLC, Shizuoka, Japan) using standard two-step collagenase perfusion method.^[25]^ Isolated rat hepatocytes with a viability of more than 85% as determined by trypan blue exclusion test were used.

### Cell fiber formation to encapsulate primary rat hepatocytes

The cell fibers were formed by using the double-coaxial laminar-flow microfluidic device assembled from pulled glass capillary tubes (inner diameter: 0.6 mm; pulled tip diameter: ∼230 µm; outer diameter: 1 mm, G-1, Narishige, Tokyo, Japan), rectangular glass tubes and connectors as previously reported ^[16]^. To form the core-shell microfibers to encapsulate primary rat hepatocytes, three types of solutions were prepared: (1) primary rat hepatocytes-containing pre-gel solution of a mixture of two types of collagen, bovine type I collagen (AteloCell™, IAC-50, Native collagen, KOKEN, Japan) supplemented with 10 % of Matrigel*®* (Corning, Tokyo, Japan) as ECM for the core; (2) pre-gel solution of 1.0% Na-alginate (from 80 to 120 cP; Wako Pure Chemical Industries, Japan) in saline for the shell; (3) mixture of 100 mM CaCl_2_ and 3% *w/w* sucrose solution for the sheath stream. To form the core-shell hydrogel microfibers with total diameters of 154±28 µm and core diameters of 77±12 µm, the flow rates of the streams in the three layers were corresponded and adjusted as follows: the flow rate of the stream in the core, the shell, the sheath was 20 µL min^-1^, 80 µL min^-1^, and 3.6 mL min^-1^, respectively.

### Cell counting and viability assay

Counting primary rat hepatocytes and evaluating their viability per cell fiber were performed three times and averaged. Prior to those procedures, the alginate shells were removed, and the primary rat hepatocytes were retrieved. To remove the shells of the cell fibers, 4 mg mL^-1^ of alginate lyase (Sigma Aldrich) in Dulbecco’s Phosphate-Buffered Saline (DPBS) (+) was added at a 1:100 ratio to the culture media, and the media were incubated for 15 minutes. To retrieve primary rat hepatocytes from ECM, 4 mg mL^-1^ of collagenase (Sigma-Aldrich) in DPBS (+) was added at a 1:50 ratio to the culture media and the media were incubated for 5 minutes. The cell number of primary rat hepatocytes per cell fiber was counted by using cell-counting plate (WakenBtech, Japan). The cellular viabilities were examined by the trypan blue dye-exclusion test.

### Measurement of albumin and urea levels

The amount of albumin in culture medium was measured using an enzyme-linked immunosorbent assay kit (rat albumin ELISA kit, Bethyl Laboratories, Montgomery, TX). The amount of urea in culture medium was measured using an assay kit (QuantiChrom urea assay kit, BioAssay Systems, Hayward, CA). For both assays, respective absorbency at 450 nm or at 430 nm was measured using a microplate reader (MTP-810, Corona Electric, and Hitachi, Japan). All the values were normalized by the number of hepatocytes and the data were expressed as protein amount 10^−6^ cells time^-1^

### CYP assay

Activity of cytochrome P450 1A1 enzyme was assessed using ethoxyresorufin-O-deethylase (EROD) assay as described previously. ^[32]^ Briefly, 48 hours before EROD assay, primary rat hepatocytes cultured either in cell fibers or in collagen-coated well-plates started to be treated with 3 mM of the CYP1A1 inducer, 3 methylcholanthrene (3-MC; Sigma-Aldrich), contained in DMEM with a final concentration of 0.1% of dimethylsulfoxide (DMSO; Sigma-Aldrich). The media containing inducers were changed daily during the treatment. In the procedures of EROD assay, inducer-treated primary rat hepatocytes were incubated with 25 µM of EROD substrate for 1 hour and subsequently the absorbency of samples at 530 nm was measured by a microplate reader (MTP-810, Corona Electric, Japan).

### Flow cytometric analysis

Using flow cytometry, primary rat hepatocytes in cell fibers were identified by detecting Asialoglycoprotein receptor-1 (ASGPR-1) based on previous report.^[33]^ First, the hydrogel shells of the cell fibers are dissolved using alginate lyase (Sigma Aldrich), and the cells of the core were dissociated using collagenase (Brightase, Nippi, Japan). These cells were resuspended in DMEM supplemented with 10% of FBS, incubated for at least 30 minutes, and then centrifuged at 400g. The cells were treated for 30 minutes with a 100 µL solution containing a mouse-anti-human ASGPR1 antibody (Thermofisher, Waltham, MA) diluted 1:50. A fluorescein isothiocyanate (FITC)–labeled goat anti-mouse antibody (Thermofisher, Waltham, MA) was added to the solution at the concentration of 0.5 g mL^-1^, and the cells in the solution was incubated for 30 minutes. The cells were washed 3 times with 1 mL fluorescence-activated cell sorter buffer (1% sodium azide and 2% fetal bovine serum in phosphate-buffered saline), resuspended in 500 µL fluorescence-activated cell sorter buffer, and analyzed by flow cytometry. Cell analysis was performed on a Becton–Dickinson FACSVerse flow cytometer and data acquisition and analysis was performed using Becton–Dickinson DiVa software (Becton–Dickinson, San Jose, CA)

### Histological analysis

Cell fibers containing primary rat hepatocytes were fixed in 4% paraformaldehyde in a 0.1 M phosphate buffer at pH 7.4. All samples were immersed in the fixative individually for 6 h. The samples were dehydrated through graded series of ethanol, immersed into xylene, and embedded in paraffin according to the standard methodology. Then, 5-6 µm -thick sections were obtained. For H&E staining, the sections were de-waxed, underwent hematoxylin and eosin. For immunohistochemistry, the sections were de-waxed and were incubated in blocking buffer (1% goat serum in PBS). To detect albumin, rabbit anti-rat albumin antibody (RaRa/ALB/7S, Nordic-MUbio, Netherlands) was used as primary antibody and biotinylated goat anti-rabbit antibody (VECTASTAIN Elite ABC HRP Kit, Vector Laboratories, CA) were used as secondary antibody. To visualize the antigen-antibody reaction, the sections were incubation in a 0.05 M Tris-HCl buffer (pH 7.6) containing 0.01% 3,3’-diaminobenzidine and 0.001% H_2_O_2_. The sections were observed under a light microscope (BX 53; Olympus corporation, Tokyo, Japan).

### Hepatotoxicity assessment

Acetaminophen (Sigma-Aldrich) and diclofenac (Sigma-Aldrich) were adopted as hepatotoxic compounds and stocked in DMSO at the concentration of 10 mM and 1mM, respectively. Primary rat hepatocytes cultured either in cell fibers or in collagen-coated 24-well plates were treated for 24 hours with either of these compounds diluted in media at intended concentrations, and then the respective compound-treated hepatocytes underwent assays of their different specific characteristics: cell viability, albumin secretion, or urea synthesis. Data from drug-treated hepatocytes were normalized to the respective untreated controls in the same culture system. By plotting these normalized data on y-axis and the concentrations of compounds on x-axis, the concentration-reaction curves were generated. Afterwards, using Origin software (Origin 8.1, OriginLab Inc., Massachusetts), IC50 values were estimated as the interpolated concentrations at which 50% of the hepatocytes are supposed to have lost their characteristics.

### Assessment of hepatocyte proliferation inhibition

Retrorsine (Sigma-Aldrich) was adopted as a compound that inhibits hepatocyte proliferation. The working solution of retrorsine was prepared through the following two steps: 1) retrorsine was added into distilled water at 20 mg ml^-1^ and dissolved completely through titration of the solution to pH 2.5 using 1N HCl; 2) this solution was neutralized using 1N NaOH, and NaCl was added into this solution at 150 mM. Then immediately, this working solution was diluted in media at intended concentrations. The changeover between starting and stopping exposure to retrorsine was done by medium replacement; the culture media surrounding the cell fibers can be completely removed from petri dishes to be replaced with new ones while keeping the cell fibers intact. The proliferation rate of the hepatocytes in the experimental groups was calculated by dividing the numbers of the hepatocytes that proliferated from 2 days of culture to 3 days of culture in the experimental groups with the ones in the control group. The concentration-reaction curves were generated by plotting these proliferation rates on the y-axis and the concentrations of retrorsine on x-axis. Thereafter, using Origin software (Origin 8.1, OriginLab Inc., MA, USA), IC50 value for retrorsine was then estimated as the interpolated concentrations at which 50% of the hepatocytes are supposed to have been inhibited in proliferation.

### Cell fiber transplantation for NARs

Six-week-old male NARs were purchased from Japan SLC (Shizuoka, Japan). The rats were conditioned for the experiments by being housed in plastic cages in a room at the controlled temperature of 23 ± 2°C. The rats weighing more than 240 g underwent surgical procedures for transplantation of the cell fibers as follows: the rat was anesthetized by 2.0% isoflurane (isoflurane for Animal, Intervet, Tokyo) delivered using an animal anesthetizer device (MK-AT210D, Muromachi Kikai, Tokyo). Cell fibers encapsulating 2×10^7^ primary rat hepatocytes that had been cultured for 7 days *via* their proliferation under 3T3CM were rinsed with serum-free culture medium once. The cell fibers were then sucked into a 23 G butterfly needle (SURFLO, Terumo, Tokyo) connected with a 20 mL syringe. After laparotomy of the rat, small and large intestines were exposed, and the cell fibers were injected into intramesenteric space and placed along the portal vein. The rat received subcutaneous injection of tacrolimus, FK-506 (Prograf, Astellas Pharma, Tokyo) at 1 mg kg^-1^ body weight every day after transplantation. Thirty-five days after transplantation, the rat underwent second-look laparotomy under the same anesthesia used for the transplantation procedure, and the transplanted cell fibers encapsulating primary rat hepatocytes were removed. 350 μL whole blood were sampled from the rat through its tail vein before and every 7 days up to 35 days after transplantation, and the plasma albumin concentrations were measured using ELISA assay (Rat Albumin ELISA Quantitation Set, Bethyl Laboratories, Inc, TX).

### Statistical analysis

Data were processed with Origin software (Origin Lab Inc). Each experiment was carried out with at least three individual samples. All the values in results were given as the mean ± standard deviation (SD). Statistical analysis was performed by using Student’s t-test, with *P* =0.05 or P= 0.01 considered to be statistically significant, if not stated differently. For *in vitro* detection of drug hepatotoxicity (Table S3), we additionally assessed for the significance of the differences between groups by using one-way analysis of variance (ANOVA).

*More information is available in supplemental methods in supplementary information*.

## Acknowledgments

We thank S. Nagata and T. Watanabe for having productive discussions with us and K. Ikeda for his assistance for cell culture as well as for his advice on device fabrication.

We also thank K. Nara for his technical assistance with microfiber fabrication and cell preparation

This work was partially supported by Grant-in-Aid for Scientific Research (S) (Grant number: 16H06329) and was also partially supported by JSPS KAKENHI (Grant Number 19H04439).

E.M-A. is supported by JSPS Postdoctoral Fellowship for Foreign Researchers (Long-term) and was partially supported by Japan Agency for Medical Research and Development (AMED), Research Center Network for Realization of Regenerative Medicine, Projects for Technological Development.

## Authors contribution

E.M-A. T.O. and S.T. conceived and designed the experiments; E.M-A. H.T. H.A. M.K. and M.Y. performed the experimental work and contributed to the manipulation of materials, reagents, and analysis tools; T.O. M.K. H.A. and M.Y. performed animal experiments; E.M-A. T.O. and S.T. analysed the data and wrote the manuscript. All authors read and approved the final manuscript.

## Additional information

The authors declare competing financial interests: TO and ST are inventors on intellectual property rights related to the cell fiber technology, and stockholders of Cellfiber Inc, a start-up company based on the cell fiber technology.

